# Validation and classification of RNA binding proteins identified by mRNA interactome capture

**DOI:** 10.1101/2021.02.02.429302

**Authors:** Vaishali, Lyudmila Dimitrova-Paternoga, Kevin Haubrich, Mai Sun, Anne Ephrussi, Janosch Hennig

## Abstract

RNA binding proteins (RBPs) take part in all steps of the RNA life cycle and are often essential for cell viability. Most RBPs have a modular organization and comprise a set of canonical RNA binding domains. However, in recent years a number of high-throughput mRNA interactome studies on yeast, mammalian cell lines and whole organisms have uncovered a multitude of novel mRNA interacting proteins that lack classical RNA binding domains. Whereas a few have been confirmed to be direct and functionally relevant RNA binders, biochemical and functional validation of RNA binding of most others is lacking. In this study, we employed a combination of NMR spectroscopy and biochemical studies to test the RNA binding properties of six putative RNA binding proteins. Half of the analysed proteins showed no interaction, whereas the other half displayed weak chemical shift perturbations upon titration with RNA. One of the candidates we found to interact weakly with RNA *in vitro* is *Drosophila melanogaster* End binding protein 1 (EB1), a master regulator of microtubule plus-end dynamics. Further analysis showed that EB1’s RNA binding occurs on the same surface as that with which EB1 interacts with microtubules. RNA immunoprecipitation and colocalization experiments suggest that EB1 is a rather non-specific, opportunistic RNA binder. Our data suggest that care should be taken when embarking on an RNA binding study involving these unconventional, novel RBPs, and we recommend initial and simple *in vitro* RNA binding experiments.

## Introduction

Ribonucleoprotein particles (RNPs) are RNA and protein assemblies that carry out or regulate essential functions in cells, including transcription, splicing, translation and RNA decay among others (Cech and Steitz 2014). The proteins that bind RNA molecules directly – so-called RNA binding proteins (RBPs) – typically have a modular organization and comprise a set of globular RNA binding domains (RBDs). Among the most abundant RBDs are the RRM (RNA recognition motif) domains, which are present in roughly two thirds of all studied mRNA binding proteins (mRBPs), followed by DEAD box helicase domains, zinc fingers, KH domains and cold shock domains (CSDs) (Lunde et al. 2007; Gerstberger et al. 2014; Corley et al. 2020). Besides the well-defined globular domains, some RNPs consist partially or entirely of low complexity (LC) sequences including Arg-Gly-Gly (RGG) and Arg-Ser (RS) repeats as well as positive Lys/Arg (K/R) patches (Balcerak, 2019). Many of these LC proteins can phase separate and are a component of membrane-less RNP granules, where they serve as platforms for protein-protein or protein-RNA interactions (Chong et al. 2018). Whereas most often RGG and RS repeats engage in low affinity, non-specific interactions, there are examples of high-affinity interactions and co-folding with target RNAs (Phan et al. 2011). Other examples of non-canonical RBD interactions can be found within large RNPs such as pre-ribosomal particles, ribosomes and spliceosomes. With the recent X-ray and cryo-EM structures of the eukaryotic ribosome, it has become clear that, in contrast to their prokaryotic counterparts, many eukaryotic ribosomal proteins have long insertions that are either unstructured or form extended helices which make contacts with the ribosomal RNA (Ben-Shem et al. 2011; Klinge et al. 2011; Klinge et al. 2012).

Earlier, RBPs were identified using biochemical methods such as UV crosslinking, followed by RNA affinity purification and identification of the bound proteins by immunoblotting or mass spectrometry (Dreyfuss et al. 1984; Pinol-Roma et al. 1988; Gerstberger et al. 2014). With the advent of whole-genome sequencing, and the determination and deposition of high-resolution structures in the protein data bank (PDB), new candidate RNA binding proteins were put forward through multiple sequence alignment and computational predictions (Gerstberger et al. 2014). During the last decade, a variety of *in vitro* and *in vivo* approaches have been developed to identify the complete set of RNA binding proteins (Castello et al. 2016b; Ryder 2016; Perez-Perri et al. 2018). To date, such RNA interactome capture approaches have been performed in diverse cell types, tissues and organisms (Baltz et al. 2012; Castello et al. 2012; Mitchell et al. 2013; Castello et al. 2016b; Ryder 2016; Hentze et al. 2018). In essence, these methods involve *in vivo* UV crosslinking of RBPs to RNA, followed by oligo-dT pulldown under denaturing conditions to isolate poly-adenylated RNA species, and mass spectrometry (MS) in order to identify the crosslinked proteins (Dimitrova-Paternoga et al. 2020). Overall, these studies demonstrated that the number of RNA bound proteins can reach up to 10% of the organism’s proteome in some species (Gerstberger et al. 2014; Hentze et al. 2018). Moreover, about half of the identified proteins lacked classical RNA binding domains and many did not even have previously known functions related to RNA (Hentze et al. 2018). Subsequently, a few of these proteins were tested and validated to be *bona fide* RNA binding proteins, for example p62 or the TRIM family proteins Brain tumor and TRIM25 (Loedige et al. 2014; Loedige et al. 2015; Choudhury et al. 2017; Horos et al. 2019; Haubrich et al. 2020). Of note, a verified RNA binding protein is the cytoskeletal protein APC (Adenomatous polyposis coli). APC possesses a basic stretch that was demonstrated to bind to and promote localization of *β2B-tubulin* mRNA to the plus ends of growing microtubules (MTs) in neurons (Preitner et al. 2014).

Interestingly, another cytoskeletal protein, End-Binding Protein 1 (EB1), also known to interact with APC, was identified as a novel putative RNA binding protein in two independent mRNA interactome capture studies in *Drosophila* (Sysoev et al. 2016; Wessels et al. 2016). EB1 is an evolutionarily conserved protein that binds to the plus ends of MTs in a nucleotide dependent manner and regulates the plus end dynamics (Vaughan 2005; Nehlig et al. 2017). EB1 also plays important roles in recruiting other MT associated proteins (such as CLIP-190) to the plus end of MTs (Dzhindzhev et al. 2005). Studies in *Drosophila* S2 cells revealed that EB1 depletion causes a spectrum of MT associated defects, such as reduced microtubule dynamics, a drastic reduction in astral MTs, malformed mitotic spindles, defocused spindle poles, and mis-positioning of spindles away from the cell center (Rogers et al. 2002). Similar phenotypes were observed in mitotic spindles of *Drosophila* embryos microinjected with anti-EB1 antibodies (Rogers et al. 2002). EB1 thus appears to have a crucial role in regulation of MT dynamics, which in turn are essential for cellular processes such as cell cycle, transport and localization of RNA and proteins, vesicle transport and establishment of cell polarity, all of which rely on a proper MT network and dynamics. Owing to these important roles of EB1 in regulating MTs *in vivo*, it is of interest to determine if the protein interacts directly with RNA and, if so, to study the physiological significance of the interaction *in vivo*.

Consequently, we chose EB1 as one of six putative RBPs to be validated for their RNA binding properties in this study. The choice of the five other target RBPs was based on 1) being hits in mRNA interactome capture, 2) possible additional availability of RNA binding related data, 3) amenability for NMR spectroscopy (smaller full-length protein or RNA binding has been assigned to a smaller domain). Thus, we chose the following metabolic and regulatory enzymes: human Thioredoxin, *hs*TXN; yeast FK506-binding protein 1, *sc*FPR1, and human tripartite motif protein 25, *hs*TRIM25; an adapter protein (human Beta-1-synthrophin, *hs*SNTB1), and one other cytoskeletal protein, (*Drosophila* atypical Tropomyosin1 (*a*Tm1)). These candidates are described in more detail below.

As a primary tool, we chose nuclear magnetic resonance spectroscopy (NMR) to validate the RNA binding properties of the six putative RBPs, due to its high sensitivity to changes in the chemical environment of protein residues upon interaction with ligands (in this case RNA). As a result, even extremely weak interactions can be studied using NMR. This is of special importance as we do not know whether the RBPs of interest possess certain sequence specificity. Three of the proteins did not show any chemical shift perturbations (CSPs) upon RNA titration, whereas the other three did to varying degrees. One of the proteins for which CSPs indicated a weak interaction with the RNA tested was EB1. CSP analysis and further competition assays demonstrated that RNA interacts with EB1’s microtubule binding surface. To understand the physiological significance of this interaction we performed RNA immunoprecipitation (RIP) with GFP-tagged EB1 from *Drosophila* oocytes. Verification of some of the most enriched targets by colocalization analysis and EB1 knock-down indicate that EB1 is rather an opportunistic RNA binder. As it stands, we propose to categorize novel RBPs devoid of a classical RNA binding domain into independent, dependent and opportunistic RBPs.

## Results

### NMR titration studies of six RBPs against poly(U) oligo RNA

In order to obtain biophysical evidence of direct RNA binding, we performed NMR-monitored RNA titrations of putative RBPs lacking a canonical RNA-binding domain, selected from hits of RNA interactome studies in yeast, *Drosophila* and mammalian cell lines. We selected TXN and SNTB1, as data exist regarding their putative RNA binding surfaces and it would be relatively straightforward to confirm these surfaces by NMR. We expressed full-length proteins (*dm*EB1, *sc*FPR1, *hs*TRX) or domains, to which RNA binding has been assigned (TRIM25-PRY/SPRY domain, *hs*SNTB1’s PDZ domain, and the N-terminal domain of *Drosophila a*Tm1) in *E. coli*, using culture medium containing ^15^NH_4_Cl as the sole nitrogen source. The purified ^15^N-labelled proteins were then titrated with synthetic RNA and HSQC spectra were recorded to detect possible interactions. As the RNA targets of the proteins are unknown, we used short poly-(U) oligomers in the initial experiments. These oligomers also serve as a good starting point because the RNA interactome studies use protein-RNA crosslinking, which only occurs between protein residues and non-paired RNA bases. At each titration point we collected a ^1^H,^15^N-HSQC. This experiment resolves each amide proton/amide nitrogen correlation in the backbone and the proton-nitrogen correlation in side chains of asparagines, glutamines and tryptophans as a single cross peak. Although the spectrum itself contains no readily extractable structural information, it is sensitive to even minor changes in the conformation and chemical environment of amino acids. Upon titration, spectral changes such as chemical shift perturbations (CSPs), line broadening and resulting signal loss are therefore very sensitive indicators of even weak and transient interactions. The induced dose-dependent CSPs, corresponding to fast exchange, also allows the determination of binding affinity, and this is often observed for single classical RNA binding domains like RRMs, KH domains, or CSDs and dsRBDs (Ankush Jagtap et al. 2019; Hollmann et al. 2020)

#### Thioredoxin (TXN)

TXN is a highly conserved enzyme which catalyzes the reduction of disulfide bonds and plays a critical role in the maintenance of redox homeostasis (Lee et al. 2013). The protein was identified in human and *S. cerevisiae* mRNA interactome studies as a putative RBP (Castello et al. 2012; Beckmann et al. 2015). Moreover, the potential RNA-binding interface was suggested to involve two conserved lysines (K3 and K8) in the N-terminal region, with K8 being at the start of α-helix α1 (Castello et al. 2016b). Addition of a poly(U) 8-mer at an excess of 5 equivalents to the protein did not induce any CSPs in its ^15^N,^1^H-HSQC NMR spectrum (Figure 1A). We extended the investigation by two more titrations with poly(A) and poly(C) 8-mers to test whether TXN could have specificity towards other bases. Again, CPSs could not be observed (Supplementary Figure S1A). Although even a non-specific single-stranded RNA should induce CSPs for a weak RNA binder, to study the possibility of TXN binding to RNA in a structured context, we titrated TXN with yeast tRNA, which should provide a large variety of single-, double-stranded regions and structural features. However, we observed no changes in the spectra, even in the presence of a large excess of tRNA (Figure S1A). The absence of any shifts led us to conclude that TXN on its own in an isolated, *in vitro* context does not bind RNA.

**Figure 1.**
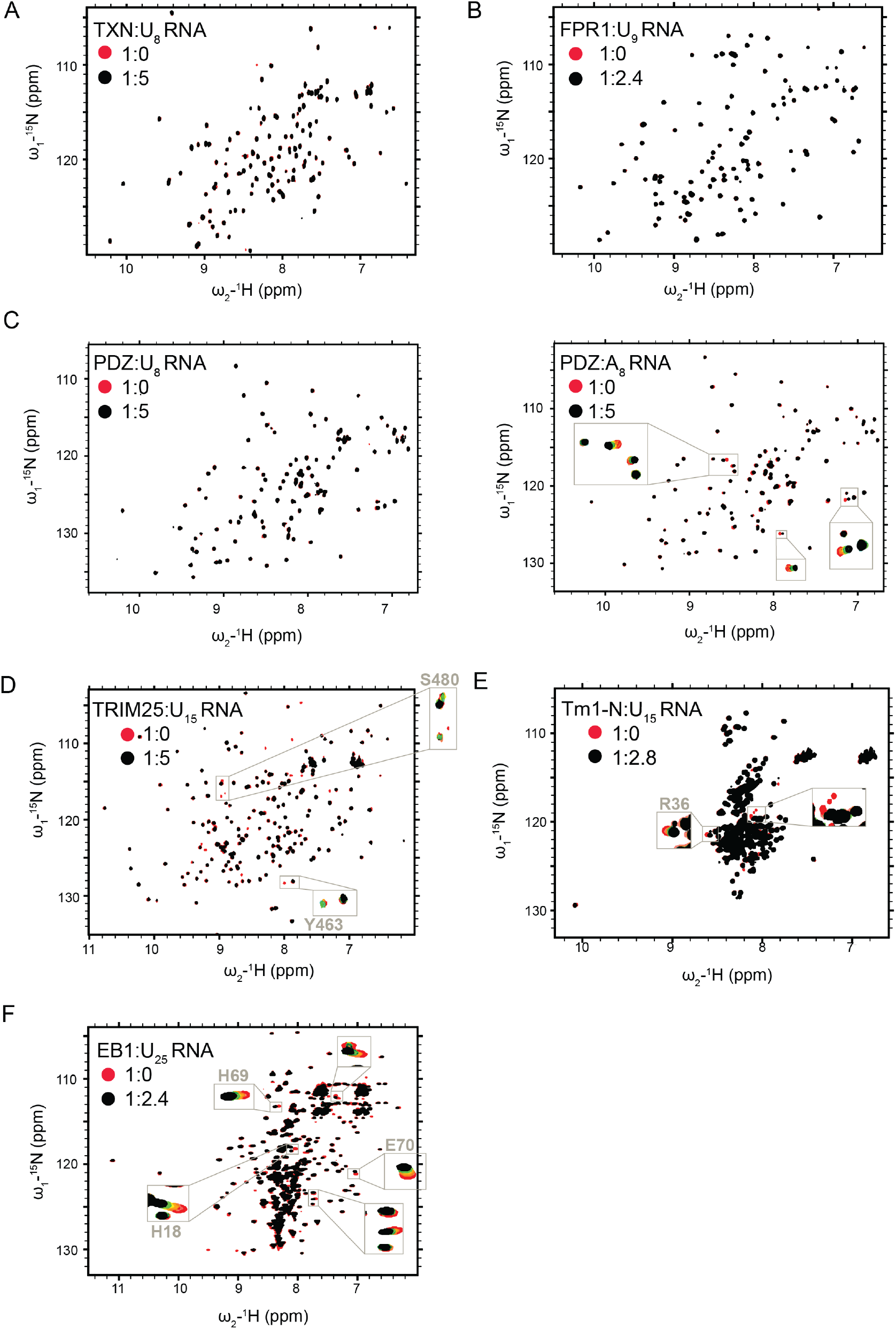
Interaction of novel RBPs with RNA: (A) ^1^H/^15^N-HSQC spectra of *hs*TXN titrated with poly(U)-8-mer, (B) *sc*Fpr1 titrated with poly(U)-9-mer, (C, left) *hsSNTB1*-PDZ domain titrated with poly(U)-8-mer, (C, right) and poly(A)-8-mer, (D) *hs*TRIM25-SPRY domain titrated with poly(U)-15-mer, (E) aTm1-N domain (I/C isoform) titrated with poly(U)-15-mer, (F) and *dm*EB1 titrated with poly(U)-25-mer. The spectra of the free proteins (in red) are overlaid with the spectra of the titrated with RNA proteins (in black). The x-axis corresponds to the ^1^H dimension whereas the y-axis corresponds to the ^15^N dimension. Each peak represents an NH bond and depicts an individual residue. The ratio of protein to RNA is also shown. For EB1, Tm1 and TRIM25, full titration points for selected residues are shown in zoomed insets (protein:RNA = 1:0/0.2/0.6/1/1.6/2.4 (EB1); 1:0/0.2/0.6/2/2.4/2.8 (Tm1); 1:0.15/0.6/1.2/2.5/5 (TRIM25); 1:0.5/1/2/3/4/5 (PDZ); 1:1/2/3/4/5 (TRX) ratios).

#### FK506-binding protein 1 (scFPR1)

*sc*FPR1 (FK506-binding protein or FKBP12 in human) is a peptidyl prolyl isomerase (PPI) identified in a yeast mRNA interactome study (Beckmann et al. 2015). The human ortholog FKBP12 is a target of the immunosuppressants FK506 and rapamycin (Hausch et al. 2013; Kolos et al. 2018). Similar to TXN, neither addition of 2.4 molar excess of a poly(U) 9-mer, nor addition of yeast tRNA at a molar excess of 5-fold induced any CSPs (Figure 1B and S1B). Of note, tRNA was suggested to be the RNA target of FPR1 based on enhanced (eCLIP) experiments (unpublished data). For comparison, we titrated rapamycin, which has been demonstrated to be a *bona fide* ligand of *sc*FPR1. Here, clear CSPs were observable, excluding the possibility that the protein used for the RNA titration experiments was inactive (Figure S1C). Thus, we conclude that *sc*FPR1 is also not an RBP in an isolated, *in vitro* context.

#### Syntrophin-beta-1 (hsSNTB1)

Syntrophins form a group of adapter proteins that link a variety of ion channels and signaling proteins to the dystrophin complex at neuro-muscular junctions (NMJ) (Belhasan and Akaaboune 2020). They feature two PH (Pleckstrin homology) domains, flanking a PDZ (PSD-95/Dlg/ZO-1) domain, and a unique C-terminal syntrophin domain (SU). Like Thioredoxin, Beta-1-syntrophin (SNTB1) and closely related SNTB2 were identified as novel RBPs in a HeLa cell mRNA interactome study (Castello et al. 2016a). Moreover, a putative RNA-binding site was mapped subsequently to a basic cavity formed by the second and third β-strands and a short helical element in between (Castello et al. 2016b). Therefore, we focused on the PDZ domain of SNTB1 for further analysis and added a poly(U) 8-mer oligomer RNA up to a 5 molar excess. Yet again, no CSPs were observed. We also tested poly(C) 8-mer for which no CSPs could be observed. For poly(A) 8-mer, however, five resonances showed small CSPs. Binding is very weak and estimated to be in the millimolar range. The magnitude of the CSP between the fourth and last titration point was as large as in the two previous titration steps and is far from saturated even at a ratio of 1:5 (protein : RNA). Thus, binding is too weak to be of physiological relevance. On the other hand, the base specificity suggests that there are more than unspecific transient charge-charge interactions, as CSPs should have been observed in this case also for poly(U) and poly(C). We conclude that the PDZ domain of SNTB1 could bind RNA in a cellular context as part of an RNP complex and falls into the category of a dependent RBP.

#### Tripartite motif protein 25 (TRIM25)

TRIM25 is a ubiquitin E3 ligase involved in cell cycle regulation, organ development and innate immunity (Orimo et al. 1999) (Zhang et al. 2015a) (Lee et al. 2018). It is a member of the TRIM protein family characterized by a tripartite motif at its N-terminus containing a RING domain, two B-Box domains and a coiled-coil region. The C-terminal region can feature a diverse set of domains in the TRIM family. TRIM25 belongs to the PRY/SPRY subfamily of TRIM proteins and thus carries a C-terminal PRY/SPRY domain (Williams et al. 2019). Besides being identified and validated as an RNA-binding protein in genome-wide screens (Kwon et al. 2013; Castello et al. 2016b), several biochemical studies *in vitro* have also reported RNA binding by TRIM25 (Manokaran et al. 2015; Choudhury et al. 2017; Choudhury and Michlewski 2019). By now, the ability of TRIM25 to bind to a number of RNAs is well established (Kwon et al. 2013; Manokaran et al. 2015; Choudhury et al. 2017; Meyerson et al. 2017; Lin et al. 2019), However, there is still some uncertainty about which part of the protein is required for RNA binding. Whereas most studies appear to confirm an important role of the PRY/SPRY domain in RNA binding (Castello et al. 2016a; Choudhury et al. 2017), the original work of Kwon et al. (2013), at least two subsequent studies also indicate direct RNA binding to the coiled-coil domain (Kwon et al. 2013; Lai et al. 2019; Haubrich et al. 2020). PRY/SPRY domains are typically protein-protein interaction domains covering a wide spectrum of substrates, ranging from linear peptide epitopes to multi-protein assemblies such as antibodies or viral capsids. The domain consists of a β-sandwich with a highly conserved core and several poorly conserved, flexible loops accommodating the substrate binding sites (Song et al. 2005). As opposed to TXN, scFPR1, and SNTB1, NMR resonances of TRIM25 exhibited clear CSPs upon addition of five molar equivalents of poly(U) 15-mer RNA, confirming its ability to bind single-stranded RNA with a dissociation constant in the high micromolar range (poly(U)-15-mer, Figure 1D, data cannot be reliably fitted due to the weak chemical shift perturbations). Interestingly, we observed binding to two distinct regions of the PRY/SPRY domain (residues 456-511 and 549-605) of which only one was previously identified by RBDmap in (Choudhury et al. 2017) (residues 470-508). In follow-up NMR studies and a detailed biophysical characterization of the RNA binding of TRIM25 that has been published in the meantime (Haubrich et al. 2020), we could show that these binding sites on the PRYSPRY domain have distinct preferences for single and double stranded RNA and thereby mediate structure specific binding to stem-loop RNAs. Additionally, we could also confirm that the coiled-coil binds to RNA synergistically with the PRY/SPRY domain. TRIM25 binding by RNA has an effect on RIG-I ubiquitination and the interferon response, suggesting an involvement of RNA in the host defense against viral infection (Haubrich et al. 2020).

#### Atypical Tropomyosin1 (aTm1)

*Drosophila melanogaster* atypical Tropomyosin1 (*a*Tm1) is a unique isoform of the actin-binding protein Tropomyosin 1 (isoform I/C), and has a role in recruitment of the motor protein Kinesin-1 to *oskar* mRNA in the *Drosophila* oocyte (Veeranan-Karmegam et al. 2016; Gáspár et al. 2017). *a*Tm1 comprises a unique, unstructured N domain, and it was suggested that the protein might directly interact with RNA (Gáspár et al. 2017). In addition, Tm1 peptides were identified in an mRNA interactome capture study from *Drosophila* embryos (Sysoev et al. 2016), although the isoform from which they originated is unknown. Upon addition of a poly(U) 15-mer RNA oligomer to the disordered N-terminal domain (aa 1-247) in a 1:2.8 ratio, we could observe CSPs and line broadening of NMR signals, clearly indicative of RNA binding (Figure 1E) with a dissociation constant of 20 μM. After backbone assignment, the largest CSP could be assigned to residue R36. The functional significance and a detailed structural and biochemical characterization have been published elsewhere (Dimitrova-Paternoga et al. 2021).

#### End binding protein 1 (EB1)

As mentioned in the introduction, EB1 was identified as a novel putative RNA binding protein in two independent mRNA interactome capture studies in *Drosophila* (Sysoev et al. 2016; Wessels et al. 2016). Structurally, EB1 is a ~33 kD protein and consists of an N-terminal Calponin Homology (CH) domain and a C-terminal EB-Homology (EBH) coiled-coil domain, connected by a linker region (Akhmanova and Steinmetz 2008) (Figure 2A). The CH domain is known to be involved in microtubule binding, and the EBH domain is crucial for EB1 homodimerisation and interaction with other proteins (Akhmanova and Steinmetz 2008). We titrated a poly(U) 25-mer RNA oligomer into full-length EB1 and observed clear CSPs in the fast exchange regime, which could be fitted to a dissociation constant in the high μM-range. Thus, EB1 could also be confirmed as an RBP *in vitro*.

**Figure 2.**
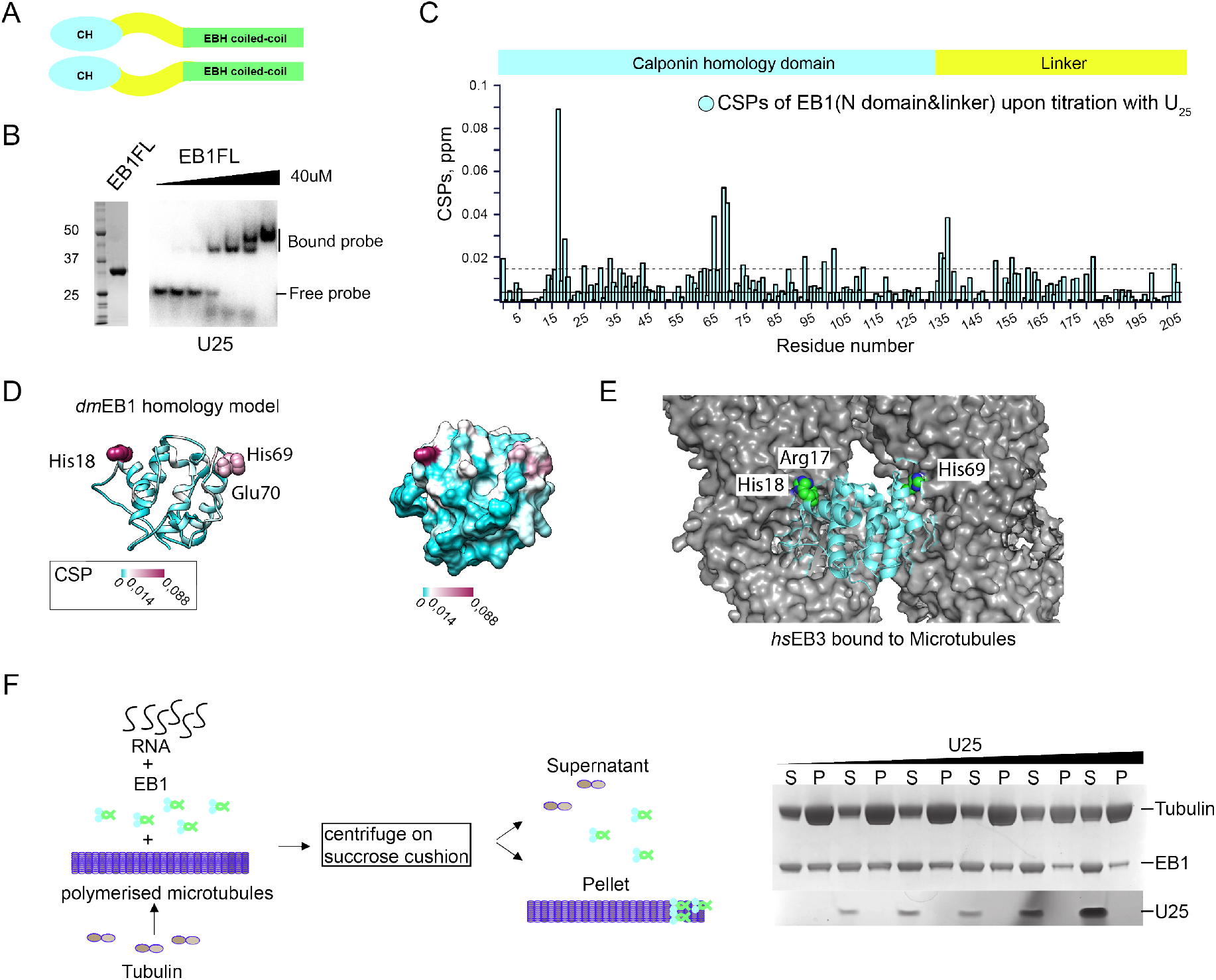
EB1 interaction with RNA involves its MT binding surface: (A) A schematic of EB1 domain organization: EB1 comprises a calponin homology domain (CH) and an EB1-homology (EBH) coiled-coil domain connected by a linker region; (B) EB1 binds RNA *in vitro*: EMSA of recombinant *dm*EB1 (left panel) with ^32^P-labelled poly(U) 25-mer (lower panel). EB1 (0,5; 1; 5; 10; 20; 40 mM protein was mixed with 2.5nM of ^32^P-labelled poly(U) 25-mer. The first lane is probe alone control ; (C) CSPs along the EB1 N+linker region upon titration with poly(U) 25-mer; (D) CSPs are plotted on the homology model of the *dm*EB1N domain in ribbon (left panel) and surface (right panel) representations. (E) Interaction of EB1 with RNA maps to the same surface with which EB1 interacts with MTs: *hs*EB3 bound to microtubules based on 3JAL (Zheng et al., 2015); (F) RNA competes with MTs for binding to EB1: Co-sedimentation of EB1 with MTs and RNA. EB1 alone or preincubated with increasing concentrations of poly(U) 25-mer were added to polymerized microtubules (left panel). After incubation, ultracentrifugation over a sucrose cushion was performed, followed by separation of the pellet and supernatant fractions. The proteins and RNA were separated on SDS PAGE and on urea denaturing gel, respectively (right panel).

In summary, half of the proteins tested could not be confirmed as independent RBPs in an isolated context, whereas the other half did show RNA binding *in vitro*. Of special interest was the confirmed RNA binding of EB1, as this protein shows the strongest CSPs, though the interaction was rather weak, as indicated by the fact that the CSPs were not saturated even at an excess of 2.4 molar equivalents of RNA. Therefore, we decided to further investigate RNA binding by EB1 in order to gain insight into its functional relevance.

### The microtubule binding surface of EB1 interacts with RNA

Titration of EB1 with poly(U) 25-mer RNA triggered CSPs in the fast exchange regime of NMR, demonstrating an interaction between the protein and RNA. We further confirmed this interaction using electrophoretic mobility shift assay (EMSA) as an additional biochemical approach *in vitro*. Adding EB1 in increasing concentrations to ^32^P-labelled poly(U) 25-mer RNA probe, gradually shifted the probe to higher molecular weight species, corroborating the interaction of EB1 with poly(U) 25-mer (Figure 2B).

In order to determine the RNA binding interface on EB1, we sought to identify the residues showing the strongest CSPs upon titration. To this end, we performed standard triple resonance backbone assignment of the calponin homology domain attached to the linker region of EB1 (Figure 2A). Because of a good overlap between the ^1^H-^15^N-HSQC spectra of EB1 full length and EB1N domain and linker region (data not shown), we could transfer the assignments to EB1 full length and plot the CSPs induced by RNA binding versus the residue number. We observed changes in three main patches – around His18 and His69 of the calponin homology domain, and around Gly138 at the beginning of the linker region (Figure 2C). Next, we plotted the changes onto a homology structure model of the CH domain of *dm*EB1 generated by Phyre2 (Kelley and Sternberg 2009). Interestingly, the CSPs map along a continuous surface that has been shown to be essential for the interaction of EB1 with MTs (Figure 2D & E, (Zhang et al. 2015b)).

The observation of a shared binding surface for RNA and MTs suggested that these might compete for the same binding site on EB1. To test this, we performed a co-sedimentation assay (adapted from (Venkei et al. 2006)) (Figure 2F), in which tubulin was polymerized into MTs in the presence of GTPγS (a non-hydrolysable analog of GTP, providing a preferential state for EB1 binding (Maurer et al. 2011)), and then incubated with a complex of EB1 and increasing concentrations of poly(U) 25-mer. Upon loading on a sucrose cushion and centrifugation, MTs accumulate in the pellet fraction together with bound EB1 and/or RNA, whereas free EB1 protein and RNA remain in the supernatant fraction (Figure 2F, schematic). Comparison of the pellet and supernatant fractions at each RNA concentration showed that with increasing quantities of RNA, the amount of EB1 pelleting with MTs decreased. This indicates that binding of EB1 to RNA and MTs is mutually exclusive (Figure 2F, right panel). As can also been seen, the quantity of microtubules that pellet down also decreases as the amount of RNA added increases, which can be attributed to our observation that EB1 appears to stimulate microtubule polymerization (Fig S2). Lack of binding of EB1 results in reduced polymerization of microtubules under the same conditions, leading to an eventual decrease of microtubules in the pelleted fraction and an increase of tubulin in the supernatant fraction (Figure S2).

### EB1 binds opportunistically to RNA *in vivo*

To understand the physiological significance of RNA binding by EB1, we aimed to identify the targets of EB1 *in vivo*. To this end, we performed an RNA-immunoprecipitation and sequencing (RIP-seq) experiment in flies expressing EB1-GFP, by pulling down UV cross-linked EB1-GFP-RNA complexes from *Drosophila* oocytes using anti-GFP antibody. RNAs so obtained were extracted and following library preparation, subjected to sequencing. This led to the enrichment of 1017 genes in the EB1-GFP sample vs GFP control with a p-value > 0.01. Out of these, based on GO term analysis for involvement in transport, localization, cell cycle and related functions, 12 candidates were selected for further validation (Figure 3A). We also analyzed the expression levels of the enriched genes and found that out of the total 1017 genes enriched, only 220 had an RPKM > 51, indicative of high expression as per the modENCODE expression level bins (Gelbart and Emmert 2013). For the candidates selected for further studies, two of these (*asp* and *chc*) are considered to be highly expressed (RPKM > 51) and two others (*drosha* and *synj*) show expression levels with RPKM values between 26 to 50. All the rest are moderately/lowly expressed.

**Figure 3.**
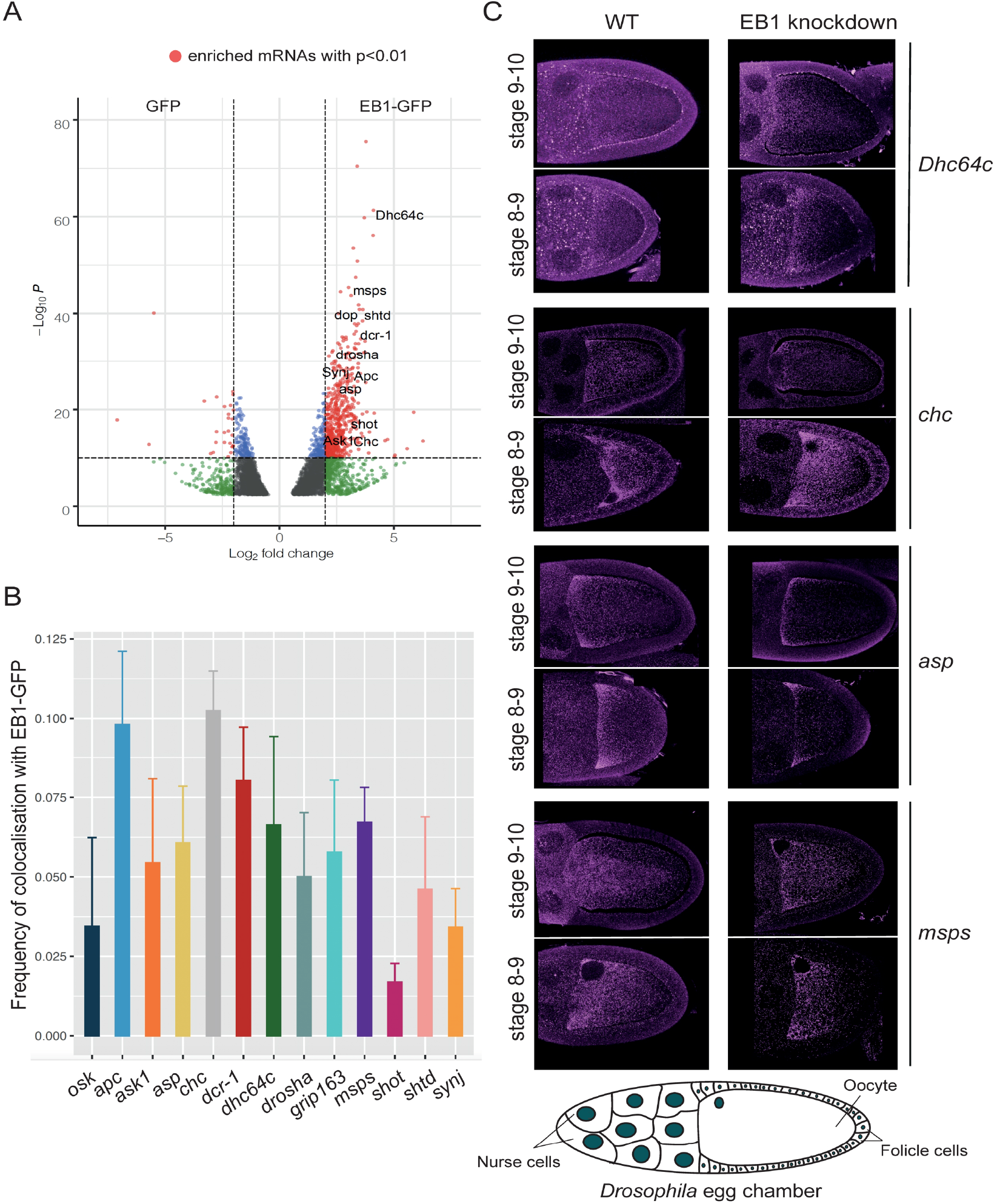
EB1 binds RNA opportunistically *in vivo*: (A) RNA immunoprecipitation-seq experiment for EB1-GFP led to the enrichment of 1017 genes with p-value <0.01: DESeq2 analysis of mRNAs pulled by EB1-GFP and GFP as a control. In red are the genes which cross the threshold of p-value<0.01 and fold change >4. (B) Frequency of colocalization of 12 of the candidate RNAs with EB1-GFP in *Drosophila* oocytes. *osk* mRNA is used as a negative control; (C) Four of the candidate genes (*chc, asp, msps, dhc64c*) exhibit a particular pattern of localization in the oocyte. Germline specific EB1 knockdown (right panels) using RNAi revealed no change in the localization pattern of these genes as compared to the wild type (left panels). A schematic of the *Drosophila* egg chamber (bottom).

To check if these target RNAs colocalize with EB1 *in vivo*, we performed single molecule fluorescence *in situ* hybridisation (*sm*FISH) of the target RNAs in oocytes of flies expressing EB1-GFP. The frequency of their colocalization with EB1-GFP was then determined. Since *oskar* mRNA, which is one of the most highly expressed RNAs in the *Drosophila* oocyte (Brown et al. 2014), was not enriched in the RIP-seq dataset, we used this mRNA as a negative control (Figure 3B). Clathrin heavy chain mRNA (*Chc*) and Adenomatous polyposis coli mRNA (*Apc*) showed the highest frequency of colocalization at 10.2% and 9.8%, respectively as compared to 5% for *oskar* mRNA, implying a relatively low frequency of colocalization of EB1-GFP with the top candidate RNA hits. We also used S2 cells, which offer a better resolution, to perform colocalization analysis of three of the top hits (*chc*, Dynein heavy chain 64c mRNA (*Dhc64c*) and mini spindles mRNA (*msps*)) with EB1-GFP.

We did not detect any colocalisation of EB1 with the candidate target RNAs in S2 cells either (Figure S2A). This implies that EB1 binds rather non-specifically to RNA and, possibly, that the RNAs which get crosslinked do so as a result of their shared subcellular localization with EB1.

Another notable feature of four of the chosen target RNAs that showed a higher frequency of colocalization with EB1 was their particular localization pattern in the oocyte. Abnormal spindle *(asp)*, *msps* and *chc* displayed an anterior localization similar to EB1-GFP, whereas *dhc64c* formed foci in the nurse cells. To see if EB1 plays a role in these localization patterns, we knocked down EB1 and assessed the distribution of the candidate RNAs. Efficiency of the knockdown in the germ line was confirmed by smFISH using *eb1*-specific probes (Figure S2 B). However, we observed no change in the distribution of any of the RNAs, suggesting that EB1 is not essential for their intracellular localization.

Taken together, these data suggest that the binding of EB1 to RNA is rather opportunistic and may not be functionally relevant. However, these experiments do not exclude the possibility that RNA might regulate the function of EB1, rather than EB1 the function of RNA.

## Discussion

In recent years extensive RNA interactome capture studies have shown that a large number of proteins without classical RNA binding domains bind directly to RNA. Here, utilizing the sensitivity of NMR to even very weak and transient interactions, we were able to test binding of several novel RNA binding proteins (RBPs) with unknown RNA targets. This has led to the identification of two main classes of RBPs. One, which can directly bind to RNAs independently of other factors (such as Tm1, EB1, TRIM-NHL (Loedige et al. 2014) and at least some TRIM-SPRY proteins), and the second, whose RNA binding ability appears to be possibly dependent on the presence of other factors *in vivo* (such as Thioredoxin (TXN), SNTB1 or FKBP12) (Figure 4). For proteins with a wide range of interactors such as TXN or SNTB1, a likely explanation is that they are a part of larger multi-protein complexes that bind RNA and are thereby brought into close proximity to RNA without actually contributing directly to RNA binding. Alternatively, additional factors present *in vivo*, but absent in our experiments could lead to allosteric changes, allowing for RNA binding of these proteins or to the establishment of joint, larger positively charged surfaces, which increase RNA affinity and specificity. This has also been demonstrated for RBPs binding RNA via classical RNA binding domains (Hennig et al. 2014; Weidmann et al. 2016). Further research should therefore concentrate on the identification of RNA binding complexes rather than RNA binding of single proteins.

**Figure 4.**
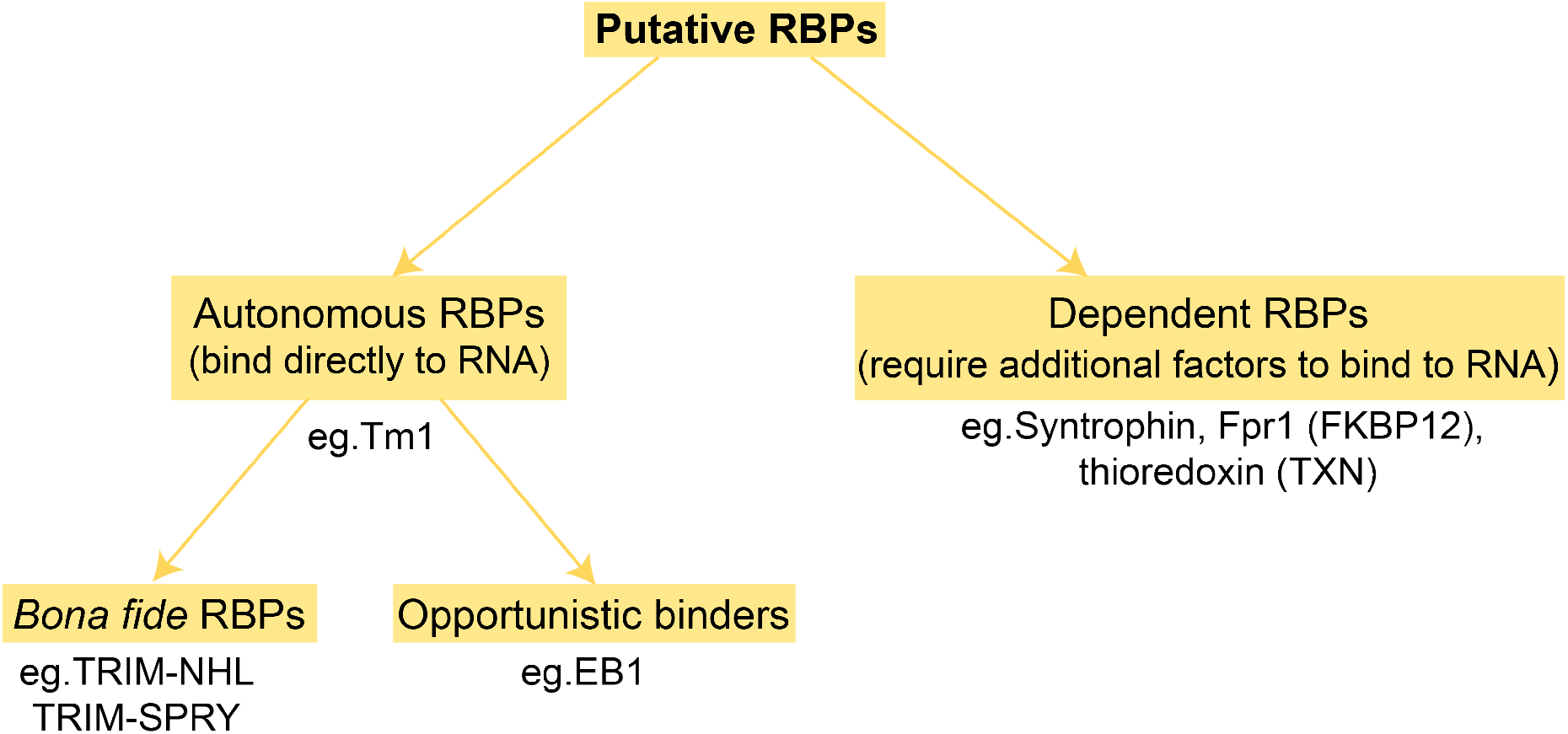
Classification of novel RNA binding proteins: Novel RBPs can be classified in two main groups: Autonomous binders such as EB1, Tm1, TRIM-NHL and at least some TRIM-SPRY proteins which bind RNA directly. The recurring validation of their RNA binding properties in individual studies changes their status from being a novel RBP to being a classical, general RBP. Others, like Thioredoxin (TXN), SNTB1 or FKBP12 however, can bind RNA only dependent on other factors present *in vivo*. It cannot be dismissed at the moment that they might not bind RNA altogether. And finally, it remains to be confirmed if proteins like EB1 which bind RNA *in vitro* do so in a functional relevant way *in vivo*.

Among the putative novel RBPs that exhibit direct RNA binding, some (Tm1-I/C, TRIM-NHL and TRIM-SPRY proteins) can be regarded as *bona fide* RBPs with their RNA binding ability confirmed both in isolation *in vitro* and *in vivo* (Haubrich et al. 2020). EB1, on the other hand exhibits RNA binding *in vitro*, but its biological significance could not be validated *in vivo*. Thus, the binding of EB1 to RNA which we observed *in vitro* might merely demonstrate electrostatic interaction with the negatively charged sugar backbone of the RNA molecule, or alternatively with its individual building blocks – the nucleotides. It was recently shown that EB1 exhibits direct binding to GTP (Gireesh et al. 2018) using the same surface it uses for binding to MTs and RNA. Other studies have also demonstrated that small GTPases or nucleotide binding enzymes are enriched in the interactome of certain RNAs (Castello et al. 2015; Liu et al. 2019). More importantly, a direct interaction of small GTPases Rab1b and ARF5 with RNA was recently demonstrated (Fernandez-Chamorro et al. 2019). Thus, proteins which bind to nucleotides might also exhibit RNA binding and likely in certain cases this interaction might have also acquired functional significance. It will be interesting to find out if the interaction of Rab1b and ARF5 with RNA also involves their nucleotide binding pocket.

Our functional analysis of the interaction between EB1 and RNA, did show that the interaction at least in the *Drosophila* oocyte appears to be rather non-specific. Firstly, the top hits from RIP-seq exhibited relatively low frequency of colocalization with EB1-GFP *in vivo*. Secondly, EB1 knockdown did not show any significant phenotypic change in the flies. All this makes it difficult to assume or decipher any meaningful interaction of the candidates with EB1 *in vivo.* However, there is a second EB1-like uncharacterized protein, CG18190, which directly interacts with EB1 (data not shown) and it can very well be that this protein compensates for the lack of EB1 when it is knocked-down in the oocyte. Further analysis will be necessary to disentangle the relationship of these two homologs and a putative role for CG18190 in RNA metabolism.

Another aspect we could not address with our experiments is the possibility of RNA regulating the function of EB1 rather than the other way around. Some recent studies have shown an active function of RNA in the regulation of certain proteins. For example, the small vault RNA regulates autophagy by controlling the oligomeric state of the p62 protein (Horos et al. 2019). In another study Huppertz et al. demonstrated that RNA can regulate the enzymatic activity of the glycolytic enzyme enolase 1 (ENO1) (Huppertz et al. 2020). Therefore, we cannot exclude the possibility that RNA might regulate the function of EB1. It is very possible that by competing with the microtubules for the same binding surface on EB1, RNA might exert an effect on cytoskeletal organization.

In summary, we showed that some novel RBPs without classical RNA binding domains do bind indeed RNA, but others do not in an isolated context. We therefore encourage and recommend a thorough *in vitro* assessment of RNA binding properties of the protein of interest before embarking on time consuming and elaborate functional studies.

## Materials and Methods

### Protein expression and purification

Full-length human TXN and SNTB1 PDZ domain (residues 111-196) were cloned into pETM11 and expressed in *E. coli* BL21(DE3). Expression was induced with 0.2 mM IPTG at OD_600_=0.6 at 18° C and cells harvested after 18h. The protein was purified by Ni-NTA affinity chromatography in 50 mM Tris, pH 7.5, 300 mM NaCl, 1 mM TCEP and eluted with a gradient of imidazole (10-300 mM). The His_6_-tag was cleaved by TEV protease and removed by a second passage over the Ni-NTA column.

Stable isotope labelled TRIM25 PRY/SPRY was expressed as previously described (Koliopoulos et al. 2018). Briefly, residues 439-630 were subcloned into pETM22 and co-expressed with KJE, ClpB and GroELS in *E. coli* BL21(DE3) (de Marco et al. 2007) in M9 media, supplemented with ^15^N-labelled ammonium chloride (Cambridge isotopes) (induction with 0.2 mM isopropyl β-D-1-thiogalactopyranoside (IPTG) at OD_600_=0.6 followed by expression at 18° C for 22 h). The protein was purified by Ni-NTA affinity chromatography and the tag was cleaved by 3C protease and removed by ion exchange. Full-length *dm*EB1 and *sc*FPR1 were cloned between NdeI and BamHI sites of pET24d-His6-Tev plasmid (see (Dimitrova et al. 2015)). *dm*Tm1 (residues 1-247) was cloned between BamHI and SacI sites of pETM11-His-SUMO plasmid (https://www.embl.de/pepcore/pepcore_services/strains_vectors/vectors/bacterial_expression_vectors/popup_bacterial_expression_vectors/). ^15^N-labelled proteins were expressed in *E. coli* BL21 (DE3) CodonPlus-RIL cells in M9 media after induction with IPTG at 18°C for 16h. Cells were lysed in 20mM Tris-HCl, pH7.5; 500mM NaCl, 0.01% NP-40, 5% glycerol, 40mM imidazole buffer supplemented with protease inhibitor cocktail (Roche) and 5mM β- mercaptoethanol. The proteins were purified by 5 ml Ni-NTA columns (GE healthcare) and eluted over an imidazole gradient (40-600 mM). After cleavage of the tag by TEV protease in the case of EB1 and FPR1 or by Senp2 in the case of Tm1-I/C, EB1 and FPR1 were further purified over Mono Q ion exchange column (GE healthcare) or Ni-NTA column for Tm1. Prior to NMR measurements, for all proteins the buffer was exchanged to 20 mM sodium phosphate, pH 6.5, 150 mM NaCl, 0.5 mM TCEP using gel filtration on a Superdex S75 16/600 or Superdex S200 16/600 column (GE healthcare).

Unlabelled EB1 for electrophoretic mobility shift assay (EMSA) or for the MT competition assay was expressed in Luria-Bertani (LB) medium.

### NMR experiments

NMR spectra were acquired at 298 K on Bruker Avance III 600 and 800 MHz spectrometers equipped with a cryogenic triple resonance probe and a Bruker Avance III 700 MHz spectrometer equipped with a room temperature triple resonance probe.

NMR titrations were done at protein concentrations of 100 μM in 20 mM Na_2_HPO_4_, 150 mM NaCl, 1 mM TCEP at pH 6.5 and 10 mM poly(U) 6-, 9- 15- or 25-mer RNA (Integrated DNA Technologies) or 2.5 mM yeast tRNA (Merck) in the same buffer were added stepwise. At each titration point a ^1^H-^15^N-HSQC was recorded. Spectra were processed using NMRPipe (Delaglio et al. 1995) and visualized using SPARKY (Goddard, T. D. & Kneller, D. G. SPARKY 3. v.3.115, https://www.cgl.ucsf.edu/home/sparky (University of California, San Francisco, 2015)). Dissociation constants from chemical shift perturbations were derived according to Fielding et al. (Fielding 2003). Backbone assignment for the EB1-Nlinker (residues 1-209) was performed using CCPNMR (Skinner et al. 2016) based HNCA, HNCACB, HNCOCACB experiments (Sattler et al. 1999). HNN and HN(C)N-correlated experiments were additionally required to assign backbone chemical shifts of *a*Tm1 due to its intrinsically disordered state (Bracken et al. 1997; Panchal et al. 2001). Experiments were recorded using apodization weighted sampling (Simon and Kostler 2019). All backbone chemical shifts have been deposited at the BMRB (EB1^1-209^ accession code: 50743, *a*Tm1^1-213^ accession code: 50940).

### Electrophoretic mobility shift assay (EMSA)

Poly(U) 25-mer RNA synthetic probe (IDT) was labelled at the 5’end with ATP, [γ-^32^P] (Hartmann Analytic) using T4 polynucleotide kinase (Thermo Fischer) and subsequently purified using Illustra Microspin G25 columns (GE healthcare). Recombinant EB1 (0.5, 1, 5, 10, 20, 40 μM) was mixed with 2.5 nM probe in 20mM Tris-HCl, 7.5; 150mM NaCl, 2mM MgCl2, 10% glycerol, 1mM DTT binding buffer in 20 ul reactions and incubated on ice for 1h. The samples were subsequently separated on 6% native 0.5X TBE polyacrylamide gel for 1h at 100V. The gel was then dried and exposed overnight to a storage phosphor screen (GE), which was finally visualized with a Typhoon Trio imager (GE healthcare).

### Co-sedimentation assay

The co-sedimentation assay was adapted from the *in vitro* polymerization assay from (Venkei et al. 2006) as follows: 60 μM porcine brain tubulin (Cytoskeleton) was polymerised into microtubules in BRB80 buffer (1x) (80 mM PIPES, 1 mM MgCl2, 1 mM EGTA, pH 6.8 with KOH) containing 2 mM GTPγS (Sigma) and 20 μM taxol (Sigma) at 37°C for 30 min. Meanwhile, EB1-RNA complexes were formed with 40 μM purified recombinant EB1 and increasing concentrations of poly(U) 25-mer RNA (0 μM to 320 μM) on ice. The complex was added to polymerised MTs and incubated for 15 minutes at room temperature. The samples were carefully layered onto 30% sucrose cushion in BRB80 (1x) buffer with taxol and centrifuged at 80,000 g for 30 minutes using a Beckman SW55Ti rotor. The supernatant was saved and the pellet was washed with BRB80 (1x) plus taxol twice, before resuspending it in BRB80 (1x) with taxol. 5 μL of the sample was loaded onto 15% urea PAGE to observe the RNA and 5 μL was loaded onto SDS-PAGE to observe tubulin and EB1, in the supernatant and pellet fractions. RNA was stained with methylene blue, and the proteins were stained with Instant blue from Expedeon.

### RNA-immunoprecipitation followed by sequencing (RIP-seq)

Ovaries were collected from flies expressing EB1-GFP, lysed in lysis buffer (20mM Hepes (pH7.5), 100mM KCl, 1mM MgCl_2_, with freshly added 80U/mL RiboLock (Thermo Scientific), 0.05% NP-40 and 1x Roche Protease Inhibitor Cocktail) and cleared at 13,200 rpm at 4 °C for 10 minutes. The lysate was cross-linked with UV at 0.3J, and subsequently was incubated with magnetic GFP trap beads from Chromotek for 1.5 hr at 4°C. The beads were then washed with high salt buffer (20 mM Hepes (pH7.5), 1M NaCl, 1mM EDTA, 0.5% NP-40 and freshly added 0.5mM DTT, 80U/mL RiboLock and 1x Roche Protease Inhibitor Cocktail), followed by medium salt (20 mM Hepes (pH7.5), 500 mM NaCl, 1mM EDTA, 0.5% NP-40 and freshly added 0.5mM DTT, 80U/mL RiboLock and 1x Roche Protease Inhibitor Cocktail) and finally low salt buffer (20 mM Hepes (pH7.5), 150 mM NaCl, 1mM EDTA, 0.5% NP-40 and freshly added 0.5mM DTT, 80U/mL RiboLock and 1x Roche Protease Inhibitor Cocktail). The beads were resuspended in 100 μL proteinase K buffer (20 mM Hepes (pH7.5), 150 mM NaCl and 1% SDS) and treated with 0.2mg/mL proteinase K (Invitrogen) for 30 min at 55°C. Following the treatment, RNA was extracted with Trizol LS, using manufacturer’s instruction. The RNAs extracted from EB1-GFP and GFP samples were used to prepare cDNA libraries using a SENSE mRNA-Seq Library Prep Kit V2 (Lexogen) and analysed by single-end 50 sequencing on an Illumina HiSeq2000. The data were analysed for differential gene expression between EB1-GFP and GFP samples using DESeq2 (Love et al. 2014).

### *Drosophila* ovaries single molecule fluorescent *in situ* hybridisation (*sm*FISH) and image analysis

Probes for the twelve candidate RNAs were labelled as in (Gaspar et al. 2017) and single molecule FISH performed as in (Gáspár et al. 2017). Two to three pairs of *Drosophila* ovaries from EB1-GFP expressing flies (Rogers et al. 2008; Sysoev et al. 2016) were dissected and fixed with 2 v/v% PFA, 0.05 v/v% Triton X-100 in PBS (pH 7.4) for 20 minutes on orbital shaker. The fixative was removed and the ovaries were washed twice with PBT (PBS + 0.1 v/v% Triton X-100, pH 7.4) for 10 min each. Ovaries were then prehybridised in 100 μL hybridisation buffer (300 mM NaCl, 30 mM sodium citrate pH 7.0, 15 v/v% ethylene carbonate, 1 mM EDTA, 50 μg/mL heparin, 100 μg/ml salmon sperm DNA, 1 v/v% Triton X-100) for 15 min at 42°C. 100μl of prewarmed probe mixture (25 nM per individual oligonucleotide in hybridisation buffer) was added to the prehybridisation mixture, and the sample was incubated for 2 h at 42°C. After hybridisation, the following washes were performed to remove excess probes: 1 ml prewarmed hybridisation buffer, 1 ml prewarmed hybridisation buffer:PBT 1:1 mixture, 1 ml prewarmed PBT for 10 min at 42°C, and finally 1 ml PBT at room temperature. Ovaries were mounted in 80 v/v% 2,2-thiodiethanol in PBS and viewed using Leica TCS SP8 confocal microscope. The images were analysed in ImageJ using particle detection and object based colocalization algorithm, as in (Gáspár et al. 2017).

### S2 cell transfection and *sm*FISH

EB1 was amplified from pET24d-His-TEV-EB1 plasmid using the primers 5’-CACCATGGCTGTAAACGTCTACTC-3’ and 5’-TTACTTGTAGAGCTCGTCCATGC-3’ and inserted into pENTR/D-TOPO (Invitrogen). Using Gateway LR Clonase II (Invitrogen), the insert was moved into the vector pAWG from the Drosophila Gateway Vector Collection (https://emb.carnegiescience.edu/drosophila-gateway-vector-collection). S2 cells were transfected with pA-EB1-GFP using Effectene Transfection Reagent (Qiagen) following the manufacturer’s protocol. 1×10^5^ of the transfected cells were seeded onto concanavalin A coated coverslips and incubated at 25°C for 1 hour. The cells were then washed once with PBS and incubated in 2% paraformaldehyde for 30 minutes at room temperature followed by washing with PBS and incubation in wash buffer (25% formamide and 2x Saline Sodium Citrate Buffer) for 5 minutes. 200 μl hybridisation buffer (25% formamide, 2x Saline Sodium Citrate Buffer, 0.02% BSA, 2 mM vanadyl ribonucleoside complex) containing *sm*FISH probes at a final concentration of 0.5 pg/nucleotide/mL was then added to the cells and they were incubated at 30°C overnight, in a humid chamber. Next, the cells were washed twice in wash buffer for 30 min each at 30°C, followed by incubation in 2.5 μg/ml DAPI in wash buffer for 10 minutes at room temperature. The coverslips were finally mounted on glass slides in Immu-Mount media (Thermo Scientific) and viewed using a Leica TCS SP8 confocal microscope.

## Contributions

V., L.D-P., K.H., A.E. and J.H. conceived the study and wrote the manuscript. M.S. purified FPR1. The rest of the experiments and data analysis were carried by V., L.D-P. and K.H.

## Acknowledgements

We thank the Genomics Core Facility at EMBL for performing the sequencing of the RIP samples. This work was supported by an EIPOD fellowship to L.D-P., co-funded by Marie Curie Actions Co-fund and EMBL MSCA-COFUND-FP <u>(664726)</u>. J.H. kindly acknowledges support via Emmy-Noether Fellowship (HE 7291). L.D-P., V. and this study were supported by the Priority Program SPP1935 (EP 37/3-1, 3-2) grant of the Deutsche Forschungsgemeinschaft (DFG) to A.E. and J.H. We gratefully acknowledge the support of the EMBL.

