## Supplemental Figures for "Validation and classification of RNA binding proteins identified by mRNA interactome capture"

### Supplemental material

#### Figure S1

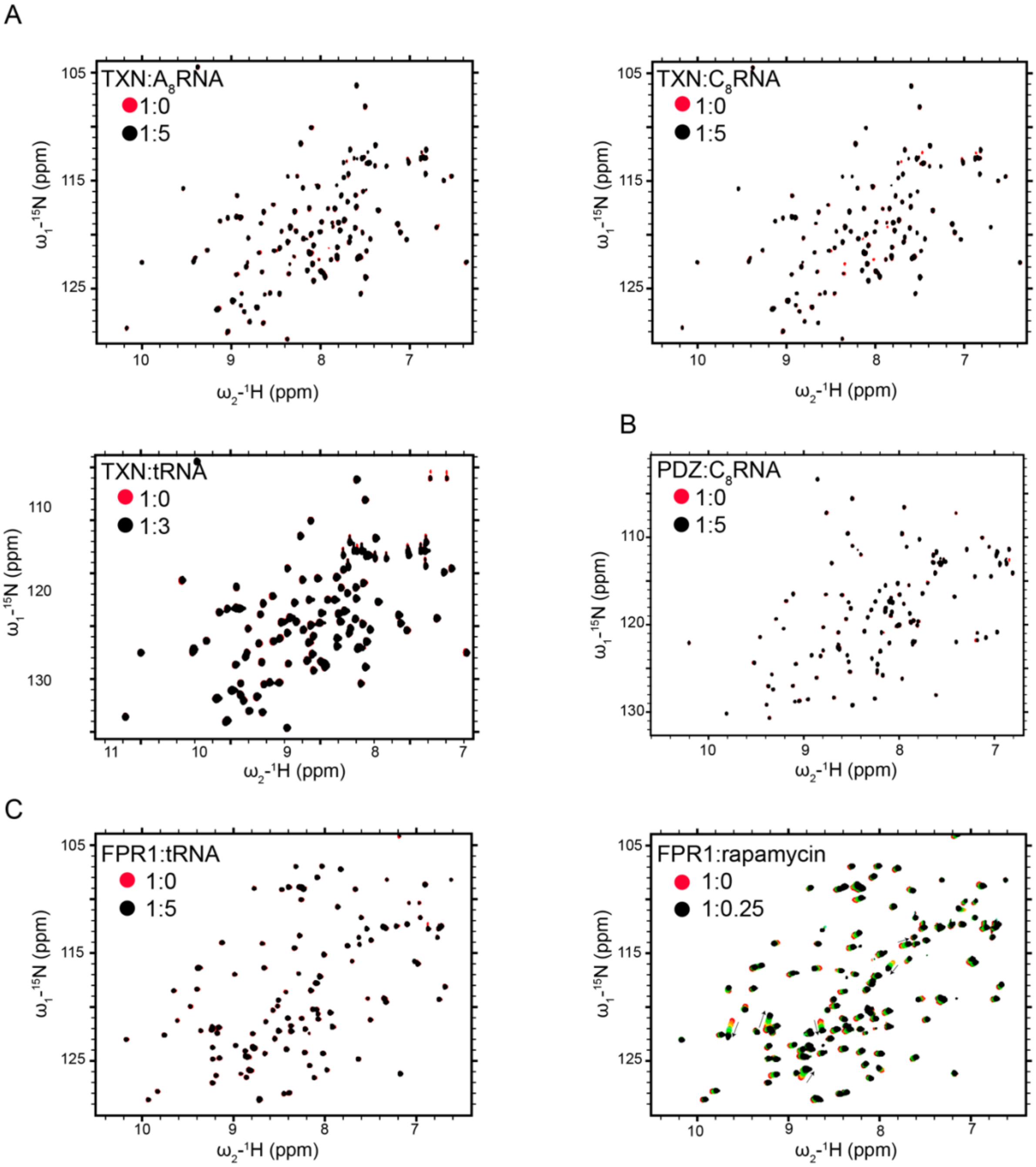


**Figure S1. (related to Figure 1)** (A) Titration of TXN with poly(A), (C)-8-mers and tRNA (no CSPs can be observed). (B) Titration of PDZ with poly(C)-8-mer (no CSPs can be observed). $^{1}H/^{15}N$-HSQC spectra of free TXN and PDZ (lower panel) (in red) overlaid with spectra of the respective proteins titrated with tRNA (in black). (C) Titrations of FPR1 with tRNA do not induce CSPs (left panel). Titration of FPR1 with rapamycin (right panel) demonstrates that FPR1 is functional: $^{1}$H/$^{15}$N-HSQC spectrum of free FPR1 (red) overlaid with spectra of EB1 titrated with rapamycin in 1:0.01 (salmon); 1:0.03 (orange); 1:0.05 (yellow); 1:0.08 (turquois); 1:0.12 (green); 1:0.25 (black) ratios.

#### Figure S2

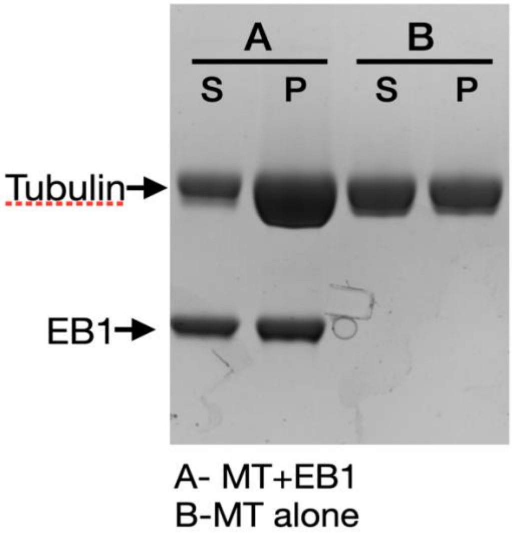


**Figure S2 (related to figure 2F). EB1 stimulates microtubule polymerization:** Tubulin was polymerized into microtubules and incubated with EB1 (A) or only buffer (B). Addition of EB1 stimulates microtubule polymerization.

#### Figure S3

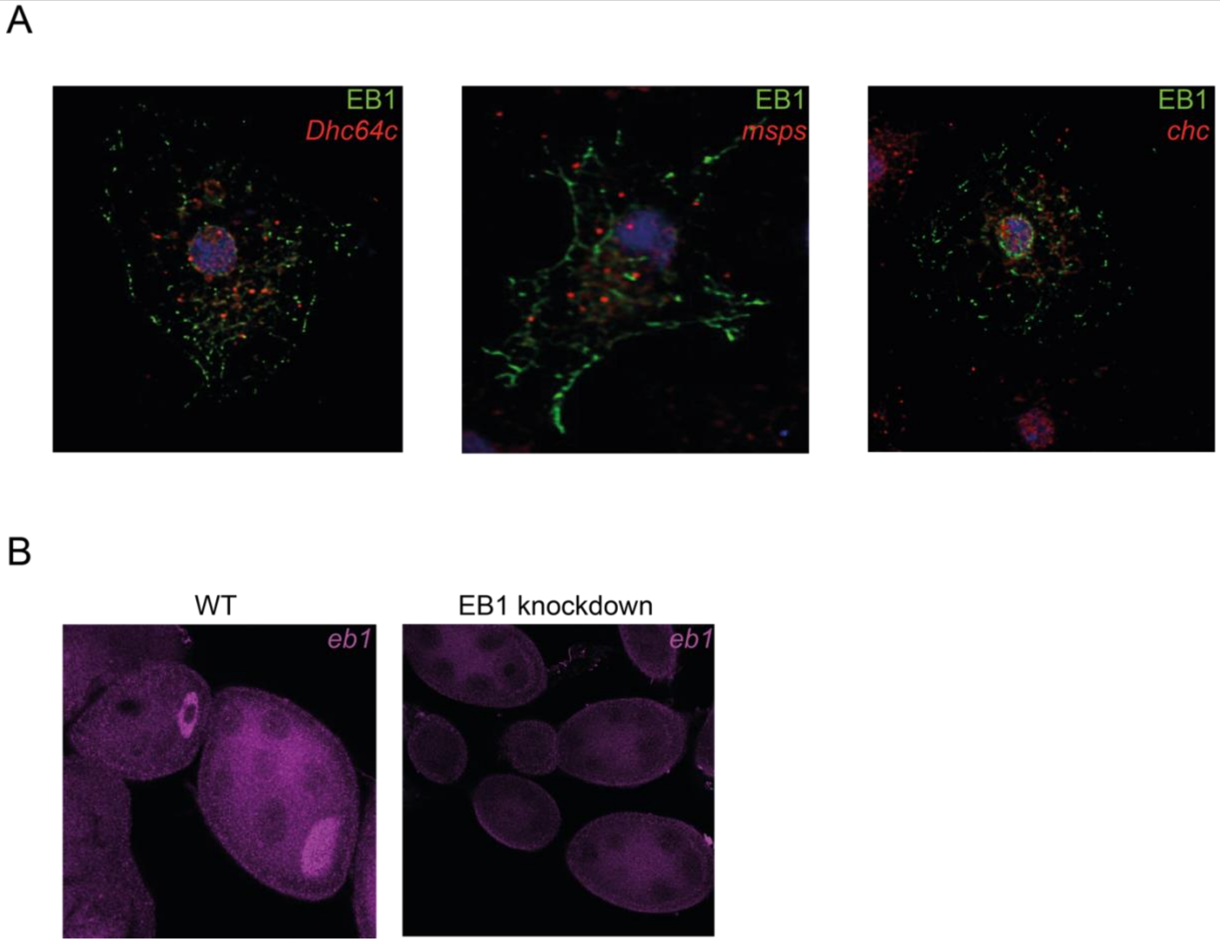


**Figure S3 (related to figure 3)** (A) Colocalization analysis of EB1-GFP and *Dhc64c*, *msps*, *chc* mRNAs. S2 cells were transfected with EB1-GFP and colocalization with the different endogenous mRNAs was detected by *sm*FISH. No colocalization was observed for any of the mRNAs; (B) EB1 was efficiently knocked down in the germ line: *sm*FISH using *eb1*-specific probes.
